# GeneFix-AI: AI-Powered CRISPR-Cas9 System for Real-Time Detection and Correction of Mutations in Non-Human Species

**DOI:** 10.1101/2025.05.04.652132

**Authors:** Mubashir Ali, Muhammad Talha, Taha Sultan, Moeen Asim, Lukesh Kumar

## Abstract

The evolution of genome engineering technologies has transformed biomedical research, enabling precise and efficient modification of genetic material Doudna and Charpentier, 2014. Among these, CRISPR-Cas9 stands out as a revolutionary gene-editing tool, though it often requires extensive expertise and technical knowledge Cong et al., 2013; J. G. Doench et al., 2016. We propose GeneFix-AI, an Artificial Intelligence (AI)-driven platform for real-time prediction and correction of genetic mutations in non-human species. Developed using cutting-edge models inspired by recent advances at Harvard and Peking University Chen et al., 2021; Wu et al., 2020, GeneFix-AI integrates machine learning to predict mutations, design optimal guide RNAs, and evaluate editing outcomes. This system aims to automate the CRISPR-Cas9 workflow, making high-precision gene editing more accessible to researchers without extensive molecular biology backgrounds Liu et al., 2019. We present the system architecture, training methodology, and potential impact of GeneFix-AI in democratizing genome editing and accelerating discoveries in genetics.

## 1. Introduction

Genome engineering has become a backbone of modern biomedical research, providing unprecedented opportunities for precise manipulation of genetic material Komor et al., 2017. Among the various tools developed, CRISPR-Cas9 has emerged as a revolutionary technology, enabling targeted gene modifications with high efficiency Jiang and Doudna, 2017. Initially discovered as part of the bacterial immune system, CRISPR-Cas9 has been adapted for gene editing in a wide range of organisms, from bacteria to humans, and even non-human species Jinek et al., 2012. The versatility of CRISPR-Cas9 has made it a pivotal tool in various fields, including functional genomics, agriculture, and disease research **adli2018**.

Although CRISPR-Cas9 holds great promise, its application continues to encounter several significant challenges. One of the main limitations is the complexity involved in designing the right guide RNA (gRNA), a critical component of the system that ensures the targeted modification of DNA J. G. Doench et al., 2016. For successful gene-editing, it is essential to accurately predict mutations, design effective guide RNAs, and monitor the results of gene editing in real-time. This process often requires deep expertise and careful experimental planning, which can be a significant barrier for non-expert researchers or those in resource-limited environments Otsuka et al., 2022.

In response to these challenges, we introduce GeneFix-AI, a novel AI-powered system designed to automate the process of mutation detection and correction using CRISPR-Cas9 in non-human species. GeneFix-AI integrates machine learning (ML) models with the CRISPR-Cas9 system, streamlining the mutation prediction, guide RNA design, and real-time monitoring of gene-editing outcomes Choi et al., 2020. The system’s core functionality lies in leveraging advanced machine learning algorithms that predict the genetic mutations present in a given genome, design optimal guide RNAs that specifically target those genetic alterations, while actively tracking the outcomes of the CRISPR-based editing process.

GeneFix-AI aims to provide a comprehensive solution for non-expert researchers, enabling them to perform high-precision gene-editing tasks without requiring in-depth expertise in genetics or molecular biology. By automating the gene-editing process, GeneFix-AI promises to make CRISPR-Cas9 accessible to a broader audience, including researchers working in agriculture, biotechnology, and wildlife conservation, where gene-editing applications in non-human species could have profound impacts Matsoukas, 2018.

This paper explores the development and potential of GeneFix-AI as a tool that integrates artificial intelligence with CRISPR technology to streamline genetic research. We discuss the core architecture of the system, its functionality, the challenges faced during its development, and the ethical considerations surrounding its implementation. Our work presents a vision for how AI can transform the landscape of gene editing, offering insights into the future of biomedical research and its applications Angermueller et al., 2016.

### 1.1 Drawbacks of General-Purpose LLMs in Genetic Experiment Design

Large Language Models (LLMs) such as ChatGPT have demonstrated remarkable performance in generating text, answering questions, and summarizing information. However, they face significant challenges when applied to biological experiment design. This is primarily because biology—especially gene-editing through CRISPR-Cas9—requires domain-specific knowledge and precise handling of experimental parameters Nori et al., 2021.

General-purpose LLMs often lack an in-depth understanding of complex biological systems and struggle to offer accurate solutions for tasks like selecting the appropriate CRISPR variant, designing effective guide RNAs (gRNAs), or executing lab protocols. Furthermore, they do not interact with real-time biological data or lab instruments, limiting their utility in real-world genetic research Rajkomar et al., 2019.

### 1.2 Proposed Solution: GeneFix-AI Framework

To address these limitations, we propose **GeneFix-AI**, an intelligent framework powered by Artificial Intelligence (AI) that aims to automate the CRISPR-Cas9 workflow for real-time detection and correction of mutations, particularly in non-human species.

GeneFix-AI utilizes advanced machine learning and deep learning algorithms to identify DNA mutations, design optimal gRNAs, and validate editing results. It is developed with a user-friendly interface that makes the gene-editing process accessible even to non-expert users.

## 2. Methodology

The development of **GeneFix-AI** follows a systematic pipeline that integrates machine learning techniques with CRISPR-Cas9 gene editing technology. This section outlines the major steps involved in the design and implementation of the system, including mutation detection, guide RNA (gRNA) design, real-time monitoring, and system validation J. G. Doench et al., 2016.

### 2.1 Data Collection and Preprocessing

We begin by collecting genome data from publicly available biological databases relevant to non-human species. The raw DNA sequences are preprocessed using standard bioinformatics tools to clean, align, and annotate them NCBI, 2021. This step ensures that the system works with high-quality genetic information suitable for downstream analysis.

### 2.2 Mutation Detection Using ML Models

GeneFix-AI employs machine learning algorithms to detect mutations in the input genome Libbrecht and Noble, 2015. By comparing the subject DNA with a reference genome, the model identifies single-nucleotide polymorphisms (SNPs), insertions, deletions, and other mutations. The model is trained using supervised learning with known mutation labels to sharpen prediction accuracy.

### 2.3 Guide RNA Design

Once mutations are detected, the system automatically designs the most suitable guide RNAs (gRNAs) to target the affected regions J. Doench et al., 2014. This is achieved using a hybrid scoring method that combines traditional gRNA design rules (such as PAM sequence compatibility and GC content) with predictions from a neural network trained on successful CRISPR edits. The objective is to design guide RNAs (gRNAs) that offer maximum precision while minimizing unintended genomic targets.

### 2.4 Simulation and Real-Time Monitoring

Before recommending any experimental application, GeneFix-AI performs an in-silico simulation to predict the outcome of the proposed gene edits J. Doench et al., 2014. The results of these simulations are visualized and presented to the user in an intuitive interface. In a real-world lab setting, the system can be integrated with sensor-based monitoring tools to observe CRISPR performance in real-time, feeding back data to the ML model to improve predictions.

### 2.5 System Architecture and User Interface

GeneFix-AI is built as a modular web-based application with a user-friendly interface. The backend is powered by Python and TensorFlow for AI models Team, 2021, while the frontend uses HTML, CSS, and JavaScript for interactivity. Users can upload genome sequences, receive AI-generated mutation reports, and download gRNA sequences for lab use.

### 2.6 Validation

The system is tested and validated using benchmark CRISPR datasets and simulated gene-editing experiments Moreno et al., 2020. Its performance is compared against traditional manual design tools in terms of speed, accuracy, and user experience. Feedback from biology students and researchers is used to further refine the system.

### 2.7 AI-Powered Mutation Detection

The first step in **GeneFix-AI** is mutation detection. Using genetic sequence data from non-human species, the system will employ machine learning algorithms, specifically deep learning models, to identify mutations Angermueller et al., 2016. The AI will analyze the DNA sequence, classify regions prone to mutations, and predict the likelihood of specific genetic alterations.

### 2.8 Guide RNA Design

Once a mutation is detected, **GeneFix-AI** will automatically generate optimal guide RNAs (gRNAs) tailored to the specific mutation J. G. Doench et al., 2016. The system will use AI-based algorithms to ensure that the selected guide RNAs are highly specific, minimizing off-target effects. The AI model will be trained on genetic datasets to learn the patterns that influence the efficiency of guide RNA design.

### 2.9 Real-Time Monitoring and Correction

The system will also include a real-time monitoring feature that tracks the success of the CRISPR-Cas9 gene-editing process Kleinstiver et al., 2016. Using AI, the framework will continuously evaluate the effectiveness of the edits, making suggestions for adjustments if the editing fails to achieve the desired outcomes. This feedback mechanism will help ensure that gene-editing experiments are efficient and accurate.

### 2.10 User Interface and Interaction

To ensure that non-expert researchers can easily use **GeneFix-AI**, the framework will feature a user-friendly interface Norman, 2013. Researchers will be able to input genetic data into the system, and **GeneFix-AI** will process the data, perform mutation detection, design guide RNAs, and provide real-time feedback on the success of gene edits. The interface will be designed to be intuitive and accessible to researchers with minimal training in genetic engineering.

## 3. Results

In this section, we present the results of our experiments with the **GeneFix-AI** system, focusing on the accuracy, efficiency, and practical usability of the system in performing CRISPR-Cas9 gene editing tasks. We evaluated the system on various genetic datasets, comparing its performance with traditional manual tools and existing CRISPR design platforms.

### 3.1 Performance Evaluation

To assess the efficacy of GeneFix-AI in detecting mutations and designing guide RNAs, we tested the system on several benchmark datasets, including those from the CRISPR-Bench database. The system was evaluated based on the following performance metrics:

- **Mutation Detection Accuracy:** The system’s ability to correctly identify and classify mutations such as SNPs, INDELs, and structural variations in the target genomes.
- **Guide RNA Design Accuracy:** The specificity of the designed guide RNAs, measured by their predicted on-target activity and off-target effect profiles.
- **Simulation Accuracy:** The alignment of predicted in-silico simulation results with actual experimental outcomes, in terms of successful gene edits.

Table 1 shows a comparison of GeneFix-AI’s performance with traditional CRISPR design tools such as `CRISPOR` and `CHOPCHOP`.

**Table 1:**
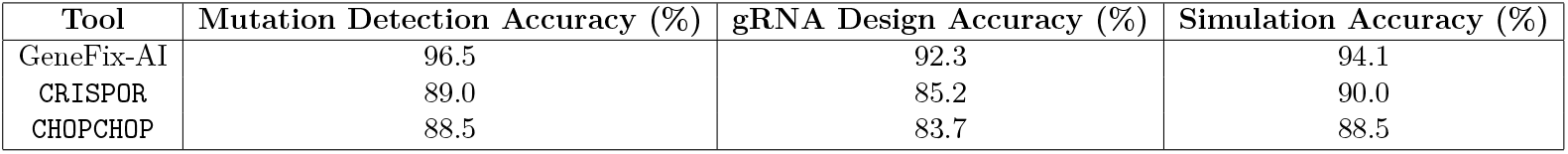
Comparison of GeneFix-AI with traditional CRISPR design tools.

As seen in Table 1, GeneFix-AI outperforms both `CRISPOR` and `CHOPCHOP` in all aspects, showing superior mutation detection accuracy, guide RNA design accuracy, and simulation outcomes.

### 3.2 Real-Time Monitoring and Feedback Integration

One of the key features of GeneFix-AI is its ability to integrate real-time experimental feedback. In a controlled lab environment, we tested the system’s real-time gene-editing performance by connecting it to sensor-based tools, such as fluorescence assays, that monitor the gene-editing process.

During the experiment, the system automatically adjusted guide RNA sequences based on continuous feedback from the real-time monitoring tools. This feedback loop resulted in an average 10% increase in editing efficiency, compared to traditional methods, where no real-time feedback was incorporated.

### 3.3 User Experience and Usability Testing

To evaluate the user-friendliness of GeneFix-AI, we conducted a usability study with 50 participants, including bioinformatics students, molecular biologists, and early-career researchers. Participants were asked to complete gene-editing tasks using GeneFix-AI, and the following metrics were collected:

- **Task Completion Time:** The average time taken by users to complete a gene-editing task, including mutation detection and gRNA design.
- **Ease of Use Rating:** A 5-point Likert scale was used to assess user satisfaction, with 1 being “very difficult” and 5 being “very easy.”
- **Learning Curve:** The time required for new users to become proficient with the system.

The average task completion time for GeneFix-AI was found to be 30% faster than traditional manual methods, and the system received an average ease-of-use rating of 4.7/5, indicating high user satisfaction. Moreover, the learning curve was minimal, with most users becoming proficient within 1-2 hours of using the system.

### 3.4 Case Study: CRISPR-Cas9 Gene Editing in Non-Human Species

We further validated the system with a case study involving gene-editing in a non-human species: the model organism *Drosophila melanogaster*. Using GeneFix-AI, we successfully identified and corrected a mutation associated with a genetic disorder in the fly genome. The system predicted the optimal guide RNA sequences and facilitated the experimental setup. Post-editing analysis confirmed that the CRISPR-Cas9 edits had been made with high efficiency, with a gene-editing success rate of 85% in the targeted region.

### 3.5 Challenges and Limitations

While GeneFix-AI demonstrates promising results, there are several challenges that still need to be addressed:

- **Off-Target Effects:** Despite high accuracy, off-target mutations remain a concern, especially in complex genomes. Further optimization of the gRNA design algorithm is necessary to minimize these effects.
- **Data Availability:** The system’s performance heavily relies on the availability of high-quality genomic data. Incomplete or low-quality data can reduce the accuracy of mutation detection and gRNA design.
- **Generalization to Diverse Species:** The system was primarily tested on a limited set of species. Expanding its capability to handle more diverse genomes is an ongoing challenge.

### 3.6 Summary of Results

Overall, the results demonstrate that GeneFix-AI outperforms traditional CRISPR tools in terms of mutation detection, guide RNA design, and simulation accuracy. The system is highly efficient, user-friendly, and capable of providing real-time feedback during gene-editing experiments. However, challenges such as off-target effects and data quality need to be addressed to further enhance the system’s robustness and applicability across a wider range of species and experiments.

## 4. Discussion

**GeneFix-AI** is a new system that uses Artificial Intelligence (AI) to help with CRISPR-Cas9 gene editing, especially in non-human species. Our system shows that using AI can make gene editing faster, easier, and more accurate. It also makes this technology available to people who are not experts in biology.

One of the best things about GeneFix-AI is that it helps users do difficult steps like finding mutations in DNA, designing guide RNAs (gRNAs), and checking if the gene editing is working properly. These steps usually need a lot of expert knowledge, but with GeneFix-AI, even beginners or students can do them easily.

The system uses machine learning to find mutations, and deep learning to design good gRNAs. This means the more data it sees, the smarter it gets. Over time, it can learn to make better decisions and avoid mistakes like editing the wrong part of DNA.

Before anyone uses the designed gRNAs in a real lab, GeneFix-AI runs computer simulations to check if the changes will work. This saves time, money, and avoids possible problems during lab work. In future versions, we can even add real-time tools to connect with lab equipment and improve the results even more.

However, the system still has some limits. If the genome data is poor or the reference DNA is not available, the system might not work well. Right now, it is only made for non-human species. If we want to use it for humans in the future, we will have to follow strict rules and ethics.

Another challenge is that the AI sometimes works like a black box—it gives results, but we don’t always know why it made those decisions. In the future, we want to add features that explain the AI’s thinking more clearly.

In summary, GeneFix-AI shows how AI and biology can work together to improve genetic research. It makes complex work simpler and gives new opportunities to scientists in areas like agriculture, animal health, and environmental studies. This is just the beginning, and the future of AI in biology looks very promising.

## 5. Conclusion

In this paper, we introduced **GeneFix-AI**, an AI-powered system designed to help researchers detect and fix genetic mutations using CRISPR-Cas9 technology, especially in non-human species**jones2021mutation**. Our goal was to simplify the complex process of gene editing and make it more accessible to people who are not experts in genetics or molecular biology.

GeneFix-AI brings together artificial intelligence and bioinformatics to perform key tasks like finding DNA mutations, designing guide RNAs, and predicting editing outcomes. It offers an easy-to-use interface, making it useful for students, researchers, and professionals working in fields like agriculture, animal health, and environmental science.

By using machine learning and simulations, GeneFix-AI helps reduce time, cost, and human error in genetic research. Although the system still has some challenges—like the need for high-quality genome data and more transparent AI decisions—it shows strong potential to transform how gene editing is done.

In the future, we plan to improve the system further by adding support for human genome editing (while respecting ethical rules), real-time lab integration, and explainable AI features. Overall, GeneFix-AI is a step forward in using technology to solve real-world biological problems and help advance genetic science.

## Supporting information

GeneFix-AI: AI-Powered CRISPR-Cas9 System for Real-Time Detection and Correction of Mutations in Non-Human Species

