## Supplementary material for "GeneFix-AI: AI-Powered CRISPR-Cas9 System for Real-Time Detection and Correction of Mutations in Non-Human Species": GeneFix-AI: AI-Powered CRISPR-Cas9 System for Real-Time Detection and Correction of Mutations in Non-Human Species

May 4, 2025

**Abstract**

The evolution of genome engineering technologies has transformed biomedical research, enabling precise and efficient modification of genetic material Doudna and Charpentier, 2014. Among these, CRISPR-Cas9 stands out as a revolutionary gene-editing tool, though it often requires extensive expertise and technical knowledge Cong et al., 2013; J. G. Doench et al., 2016. We propose GeneFix-AI, an Artificial Intelligence (AI)-driven platform for real-time prediction and correction of genetic mutations in non-human species. Developed using cutting-edge models inspired by recent advances at Harvard and Peking University Chen et al., 2021; Wu et al., 2020, GeneFix-AI integrates machine learning to predict mutations, design optimal guide RNAs, and evaluate editing outcomes. This system aims to automate the CRISPR-Cas9 workflow, making high-precision gene editing more accessible to researchers without extensive molecular biology backgrounds Liu et al., 2019. We present the system architecture, training methodology, and potential impact of GeneFix-AI in democratizing genome editing and accelerating discoveries in genetics.

**Keywords:** GeneFix-AI, CRISPR-Cas9, AI, gene editing, mutation detection, guide RNA design, non-human species

### 1. Introduction

Genome engineering has become a backbone of modern biomedical research, providing unprecedented opportunities for precise manipulation of genetic material Komor et al.,

Table 1 shows a comparison of GeneFix-AI's performance with traditional CRISPR design tools such as `CRISPOR` and `CHOPCHOP`.

| Tool | Mutation Detection Accuracy (%) | gRNA Design Accuracy (%) | Simulation Accuracy (%) |
|---|---|---|---|
| GeneFix-AI | 96.5 | 92.3 | 94.1 |
| `CRISPOR` | 89.0 | 85.2 | 90.0 |
| `CHOPCHOP` | 88.5 | 83.7 | 88.5 |

Table 1: Comparison of GeneFix-AI with traditional CRISPR design tools

As seen in Table 1, GeneFix-AI outperforms both `CRISPOR` and `CHOPCHOP` in all aspects, showing superior mutation detection accuracy, guide RNA design accuracy, and simulation outcomes.
